# The ribosomal protein RPS6A modulates auxin signalling and root development in *Arabidopsis*

**DOI:** 10.1101/2025.02.26.640312

**Authors:** Kai Pan, Kai Hou, Mengjuan Kong, Shutang Tan

**Affiliations:** University of Science and Technology of China; School of Life Sciences, Division of Life Sciences and Medicine, and Division of Molecular & Cell Biophysics, Hefei National Science Center for Physical Sciences at the Microscale, University of Science and Technology of China

**Keywords:** Ribosome, RPS6A, auxin, PIN, *Arabidopsis*

## Abstract

Protein biosynthesis by the ribosome is a fundamental biological process in living systems. Recent studies suggest that ribosomal subunits also play essential roles in cell growth and differentiation beyond their roles in protein translation. The ribosomal subunit RPS6 has been studied for more than 50 years in various organisms, but little is known about its specific roles in a certain signalling pathway. In this study, we focused on the functions of *Arabidopsis* RPS6A in auxin-related root growth and development. The *rps6a* mutant exhibited a series of auxin-deficient phenotypes, such as shortened primary root length, reduced lateral root number, and defective vasculatures. Treatment of the *rps6a* mutant with various concentrations of auxin and its analogues did not restore the root defect phenotype, suggesting a defect in the auxin signalling pathway. Further cell biological and global transcriptome analyses revealed that auxin signalling was weakened in the *rps6a* mutant and that there was a reduced abundance of PIN-FORMED (PIN) auxin transporters. Our work provides insights into the role of the protein biosynthesis pathway involved in auxin signalling.

## Introduction

Protein biosynthesis is a fundamental biological process in all living systems. The ribosome is an organelle composed of two subunits (40 S subunits and 60 S subunits), each composed of multiple components, and its central function in protein biosynthesis is highly conserved in all living lineages. The major components of ribosomes are ribosomal proteins (RPs) and ribosomal RNAs (rRNAs). The eukaryotic ribosome consists of a small 40S subunit, which plays a key role in translation initiation and contains an 18S rRNA chain and 33 proteins, and a large 60S subunit, which contains three rRNA molecules (5S, 5.8S, and 25S) and 46 protein subunits (Ben-Shem et al., 2011; Rabl et al., 2011). The ribosome not only plays an important role in protein translation but also has other functions, such as regulating the cell cycle, apoptosis, DNA repair and transcription (Lindström, 2009; Warner and McIntosh, 2009).

In the model plant *Arabidopsis thaliana*, ribosomal protein S6 (RPS6) is a component of the 40S small ribosomal subunit and contains two members, RPS6A (At4g31700) and RPS6B (At5g10360), which are functionally redundant and interchangeable (Creff et al., 2010). RPS6 is an evolutionarily conserved protein that is widely found in yeast, plants, invertebrates, and vertebrates but not in *E. coli* or archebacteria (Watson et al., 1996). RPS6 consists of an N-terminal-ribosome-bound domain (1–82 aa), a central domain (84–177 aa) and a C-terminal α-helix (178–249 aa) (Mancera-Martínez et al., 2021). RPS6 can directly cross-link to mRNAs, tRNAs, and initiation factors, determining its location in regions involved in the initiation of translation (Williams et al., 2003). There are seven phosphorylation sites of RPS6 in *Arabidopsis*, namely, Ser229, Ser231, Ser237, Ser240, Ser241, Ser247 and Tyr249, in the N-terminus, which are exposed on the surface of the 40S small subunit; thus, RPS6 can be phosphorylated by 40S ribosomal protein S6 kinase (S6K) (Obomighie et al., 2021). Studies in yeast and animals suggest that S6K-mediated phosphorylation of RPS6 is critical for its biological functions in multiple processes.

TARGET OF RAPAMYCIN (TOR) kinase is a key developmental regulator that integrates environmental and nutrient signals to regulate multiple developmental processes from embryogenesis to senescence in plants (McCready et al., 2020). S6K is a conserved phosphorylation target of TOR and regulates seedling growth by affecting the brassinosteroid (BR) signalling pathway (Dobrenel et al., 2016; McCready et al., 2020). RPS6 is downstream of the TOR kinase pathway and can be phosphorylated by S6K, which plays an important role in cellular metabolism and protein synthesis (Dobrenel et al., 2016). During active ribosome biogenesis, RPS6 is phosphorylated in the nucleolus, whereas in the cytoplasm, RPS6 is phosphorylated to promote the selective translation of a subset of transcripts (Williams et al., 2003). Therefore, phosphorylation is essential for RPS6 function.

In animals, the phosphorylation of RPS6 is involved in glucose metabolism, cell growth (Hirashita et al., 2020) and cell size regulation (Ruvinsky and Meyuhas, 2006). In *Arabidopsis*, RPS6 participates in light signalling and the auxin pathway through a TOR-dependent phosphorylation cascade, thereby regulating light-triggered translational enhancement in deetiolated *Arabidopsis* seedlings (Chen et al., 2018). The phosphorylation of RPS6 varies with photoperiod, suggesting that differential phosphorylation of RPS6A/B may contribute to the modulation of diurnal protein synthesis in plants (Blazquez et al., 2011; Enganti et al., 2017). The *rps6a* phospho-deficient mutant has a mild pointed-leaf phenotype and a shortened primary root length, which might be due to misexpression of cell cycle-related mRNAs and ribosome biogenesis proteins (Dasgupta et al., 2024). Additionally, ribosomal proteins have been reported to control developmental programs through translational regulation of AUXIN RESPONSE FACTORs (ARFs) in *Arabidopsis* (Rosado et al., 2012).

Previous studies have shown that in *rps6* mutants, the sizes of the two cotyledons are always different. The expression of many photosynthesis genes is defective, and the quantum efficiency of photosystem II is reduced in *rps6*, leading to a pale green color in the seedlings and rosette-stage plants. These effects are exacerbated at cooler temperatures (Dasgupta et al., 2024). In this study, we focused on the biological functions of the ribosomal protein RPS6A in plant growth and development. Phenotypic analysis, cell biological experiments and transcriptome sequencing revealed that RPS6A interferes with the abundance of the auxin transporter PIN and regulates auxin distribution in *Arabidopsis*. Therefore, in addition to its role in ribosomal function, RPS6A can also participate in auxin-regulated growth and development in *Arabidopsis*.

## Results

### Loss-of-function mutants of *Arabidopsis rps6a* exhibit auxin-related defects

To explore the biological function of the RPS6 protein, we generated a transfer DNA (T-DNA) insertion mutant with a loss of function in *rps6a* (Supplemental Figure 1A and B). In addition, we generated a *rps6b* CRISPR mutation in which a C-base deletion resulted in a subsequent frameshift change and premature termination of protein translation (Supplemental Figure 1C). Since both are functionally redundant and interchangeable, we used *rps6a* for physiological analysis in our study. Compared with *Arabidopsis* Columbia-0 (Col-0) wild-type (WT) plants, *rps6a* and *rps6b* mutant plants presented considerable decreases in primary root length, lateral root number and lateral root density under normal conditions (Figure 1A-E, Supplemental Figure 2E-H). In addition, the results of the root gravitropism assay revealed that, as expected, the roots of Col-0 bent toward gravity (Figure 1F and G). In contrast, the rate of root bending toward gravity was lower in the *rps6a* mutant than in the WT, indicating an impaired gravitropic response (Figure 1F and G). These phenotypic results suggest that these effects of the *rps6a* mutation on root development are reminiscent of auxin signalling dysfunction, a phytohormone that can be transported in response to gravity stimulation (Friml et al., 2003).

**Figure 1.**
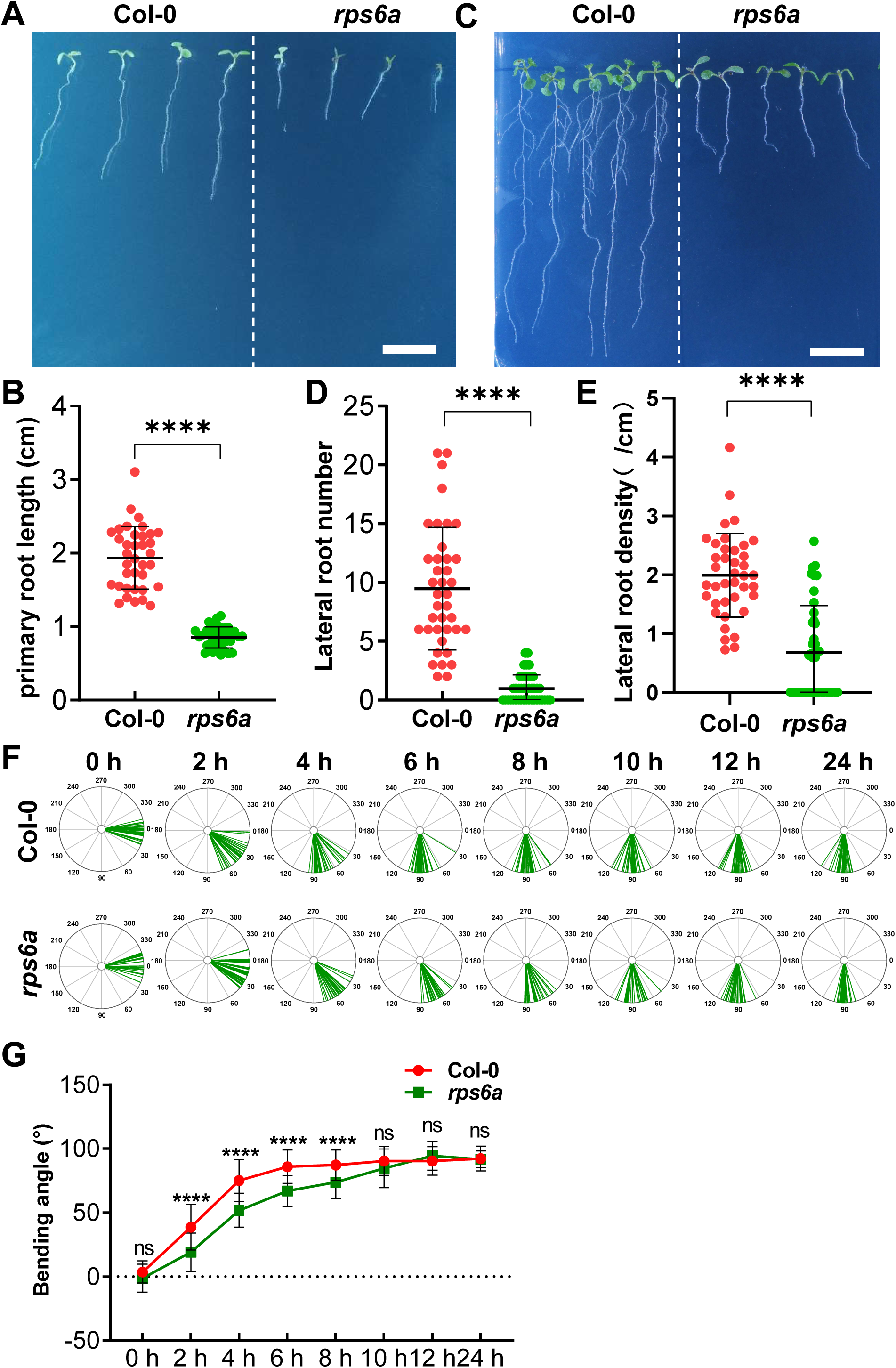
*rps6a* exhibits auxin-deficient phenotypes (A) The *rps6a* mutant has a shorter primary root. Seven-day-old seedlings were grown on MS media. Scale bar, 1 cm. (B) Quantitation of primary root lengths in Figure A, n=36, 36. The dots represent individual values, and the lines indicate the mean ± SD. \*\*\*\**p*<0.0001 according to Welch’s t test. (C) The *rps6a* mutant has fewer lateral roots. Eleven-day-old seedlings were grown on MS media; scale bar, 1 cm. (D) Quantification of the number of lateral roots in Figure C, n=39, 41. The dots represent individual values, and the lines indicate the mean ± SD. \*\*\*\**p*<0.0001 according to Welch’s t test. (E) Lateral root density in Figure C. n=39, 41. The dots represent individual values, and the lines indicate the mean ± SD. \*\*\*\**p*<0.0001 according to Welch’s t test. (F) The *rps6a* mutant has a relatively slow response to gravity. Four-day-old seedlings were rotated 90° and observed for 24 hours. (G) Quantitation of the root bending angle. At 2, 4, 6, and 8 h, the root bending angle of the *rps6a* mutant was significantly smaller than that of Col-0. The dots represent average values, and the lines indicate the mean ± SD. \*\*\*\**p*<0.0001 according to Welch’s t test.

Given that auxin has a dual role (i.e., a low concentration promotes growth, and a high concentration inhibits growth), we examined the effect of exogenously applied auxin on root growth. When grown on Murashige and Skoog (MS) media supplemented with different concentrations of auxin (indole-3-acetic acid, IAA), the auxin analogues 1-naphthaleneacetic acid (NAA) and 2,4-dichlorophenoxyacetic acid (2,4-D) (0, 50, 100, and 200 nM) (Rahman et al., 2006; Dindas et al., 2020), compared with untreated seedlings, Col-0 seedlings presented shortened primary roots (Figure 2A-F) and an increased number of lateral roots (Figure 2G-L). Notably, the *rps6a* mutant presented a considerable decrease in the number of lateral roots compared with that of Col-0 at the seedling stage (Figure 1C and D). However, the changes in primary root length and the number of lateral roots with or without chemical compounds were significantly weaker in *rps6a* than in the WT, especially in terms of the number of lateral roots (Figure 2). Additionally, when WT and *rps6a* mutant seedlings were grown on MS media in the dark, they grew upright, whereas the hypocotyls of the *rps6a* mutant were shorter than those of the WT (Supplemental Figure 2A and B). Auxin is required for leaf vasculature formation (Adamowski and Friml, 2015), and we found that the vasculature of *rps6a* mutant cotyledons was fragmented and that the leaves were smaller than those of WT cotyledons were (Supplemental Figure 2C and D). These results suggest that RPS6A impairs numerous processes involved in auxin-regulated plant growth and tropic responses.

**Figure 2.**
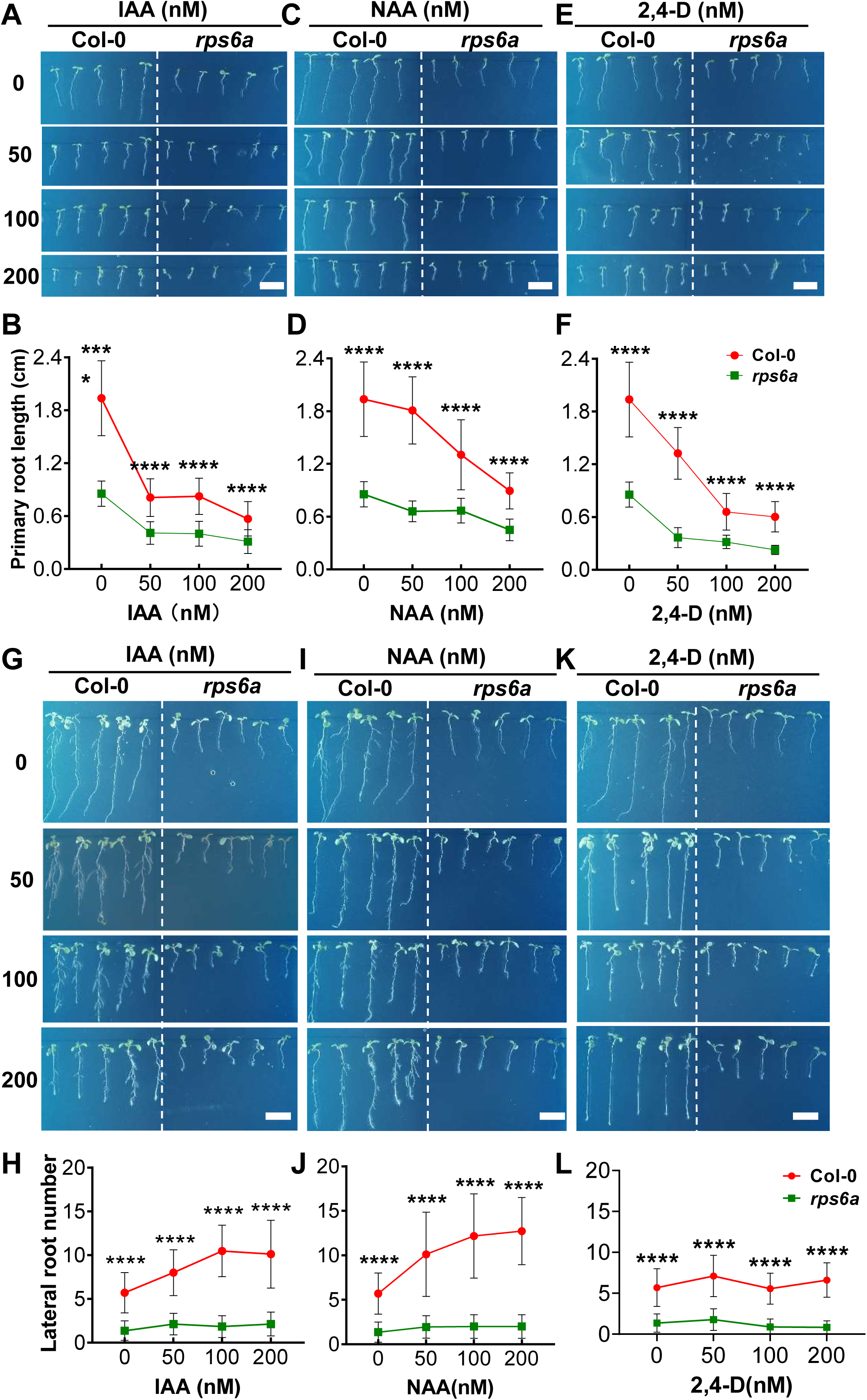
*rps6a* is less sensitive to auxin than WT (**A, C, E**) Representative images revealing that auxin and its analogues decrease primary root length. Four-day-old Col-0 seedlings were transferred to MS media supplemented with gradient concentrations of IAA/NAA/2,4-D for an additional 3 d. DMSO was used as the solvent control. Scale bar, 1 cm. (**B, D, F**) Quantitation of primary root lengths from the lines in A, C, E. (B) n=36, 18, 18, and 18 replicates for Col-0 seedlings and 36, 18, 18, and 18 replicates for *rps6a* seedlings under gradient IAA treatment. (**D**) n=36, 18, 18, and 18 replicates for Col-0 seedlings and 36, 18, 18, and 18 replicates for *rps6a* seedlings under gradient NAA treatment. (**F**) n=36, 18, 18, and 18 replicates for Col-0 seedlings and 36, 18, 19, and 18 replicates for *rps6a* seedlings under gradient 2,4-D treatment. \*\*\*\**p*<0.0001 according to Welch’s t test. (**G, I, K**) Auxin alone and its analogues did not increase the number of lateral roots of *rps6a*. Eight-day-old Col-0 seedlings were transferred to MS media supplemented with gradient concentrations of IAA/NAA/2,4-D for an additional 3 d. DMSO was used as the solvent control. Scale bar, 1 cm. (**H, J, L**) Quantitation of primary root lengths from the lines in A, C, E. (B) n=36, 18, 17, and 18 replicates for Col-0 seedlings and 36, 18, 18, and 16 replicates for *rps6a* seedlings under gradient IAA treatments. (**D**) n=36, 18, 18, and 18 replicates for Col-0 seedlings and 36, 18, 18, and 18 replicates for *rps6a* seedlings under gradient NAA treatment. (**F**) n=36, 18, 18, and 18 replicates for Col-0 seedlings and 36, 18, 18, and 18 replicates for *rps6a* seedlings under gradient NAA treatment. \*\*\*\**p*<0.0001 according to Welch’s t test.

To further investigate the relationship between auxin and root growth, we treated Col-0 and *rps6a* with different concentrations of NPA. The primary root length was significantly inhibited at 1 μM NPA (Supplemental Figure 3A and B). The root tip angle of *rps6a* was more dispersed and the phenotype is more pronounced after NPA treatment (Supplemental Figure 3C). The number of lateral roots in Col-0 did not change significantly at low NPA concentrations and decreased at higher NPA concentrations (Supplemental Figure 4). However, the lateral root number of *rps6a* was significantly inhibited after 0.1 μM NPA treatment and has no lateral roots when treated with 0.2 μM NPA, indicating that *rps6a* was more sensitive to NPA. NPA treatment aggravate the gravitropic and lateral root defects in *rps6a*, suggesting that the auxin response of *rps6a* is defective.

### Deficiency of RPS6A affects the local auxin distribution

Next, to further test whether RPS6A affects auxin distribution or signalling pathways, we examined the subcellular localization of the auxin reporter DR5rev::GFP (Friml et al., 2003). In *Arabidopsis* root tips, auxin accumulates locally via PIN proteins, which are required for root meristem activity and the gravitropic response (Adamowski and Friml, 2015; Marhava et al., 2018). Treatment with the auxin transport inhibitor NPA could lead to an enlarged DR5 expression region and overproliferation of root columella cells. A similar phenomenon was also observed for genetic mutants defective in auxin transport. For example, our previous study revealed that the apical DR5 region was enlarged in the *pdk1.1 pdk1.2* mutant, a kinase that affects PIN protein activity, which was caused by the overproliferation of root columella cells (Tan et al., 2020).

Therefore, we generated a cross between *DR5rev::GFP* and *rps6a* to analyse changes in auxin distribution or signalling. Microscopic analysis revealed that DR5rev::GFP was expressed in vascular tissues in Col-0 seedlings in the absence of IAA, with the highest expression in root columella cells (Figure 3). However, DR5rev::GFP in *the rps6a* mutant presented a broader region of columella cells without IAA treatment (Figure 3). Since exogenous application of auxin can activate the canonical transcriptional auxin pathway, resulting in increased *DR5rev::GFP* expression, we detected no change in the GFP signal after IAA treatment in *rps6a* seedlings, which is different from the findings in Col-0 (Figure 3A). The results of the GFP fluorescence signal were consistent with those of the root tips of *rps6a* seedlings, which responded to gravistimulation by 90° reorientation more slowly than Col-0 did (Figure 1F and G). Therefore, we hypothesized that RPS6A may affect auxin transport or redistribution in plants.

**Figure 3.**
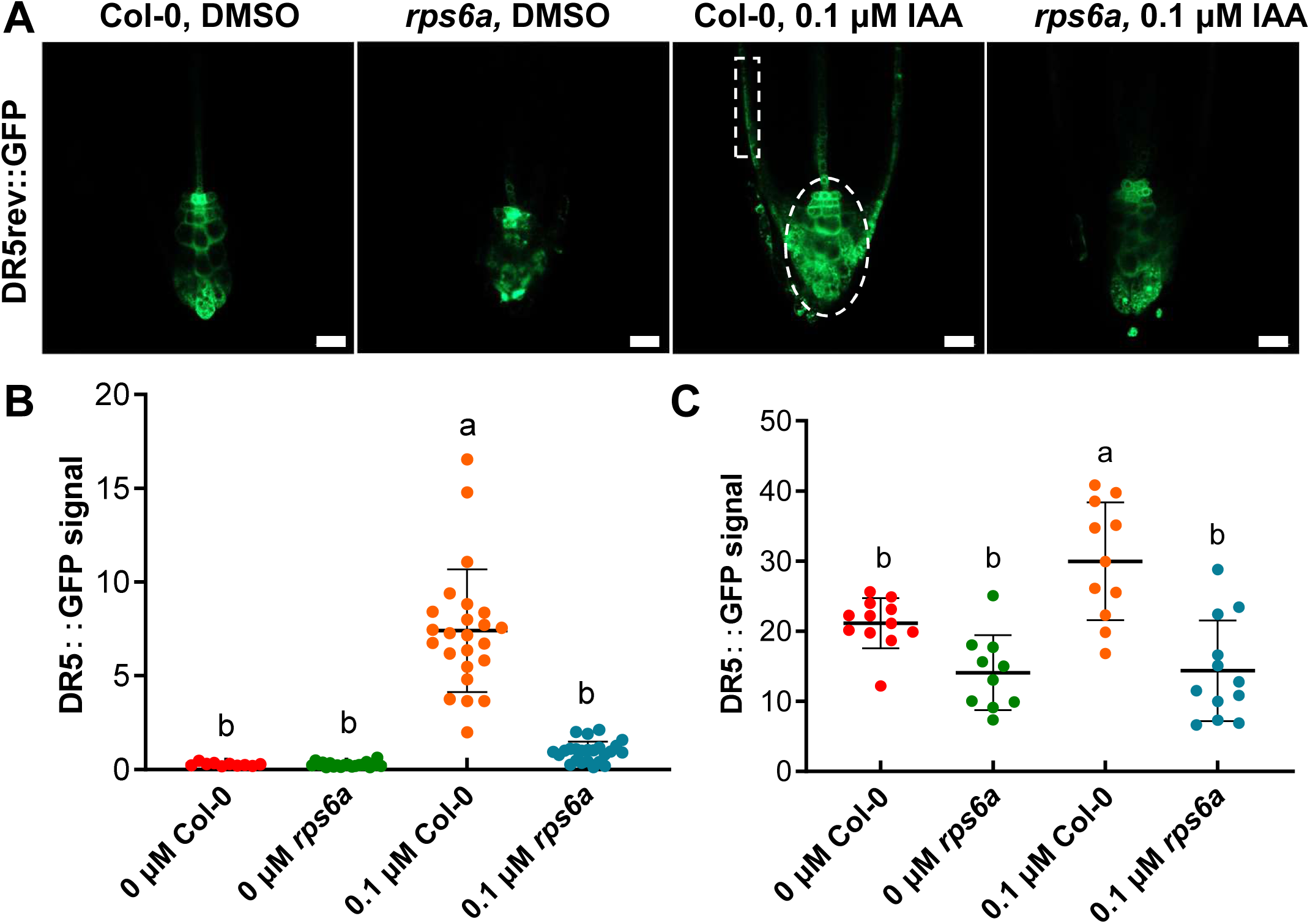
IAA treatment did not rescue the decrease in the *DR5rev::GFP* signal in the *rps6a* mutant. (A) The *DR5rev::GFP* signal decreased in the *rps6a* mutant, and the signal did not increase after 24 h of auxin treatment. Three-day-old *DR5rev::GFP* from Col-0 and *rps6a* seedlings were transferred to MS media supplemented with 0.1 μM IAA for an additional 1 d, and DMSO was used as a control. The root tips were imaged via confocal laser scanning microscopy (CLSM) in the GFP channel, 40×. Scale bar, 20 μm. (B) The GFP signal in the epidermis was measured to indicate the fluorescence intensity. n = 24, 20, 24, 24. The dots represent individual values, and the lines indicate the mean ± SD. \*\*\*\**p*<0.0001 according to Welch’s t test. (C) The GFP signal in the root tip was measured to indicate the fluorescence intensity. n = 12, 11, 13, 13. The dots represent individual values, and the lines indicate the mean ± SD. \**p*<0.05 according to Welch’s t test.

### RPS6A regulates auxin transport by affecting the abundance of PIN proteins

The above physiological phenotype and cell biology experimental results show that RPS6A might affect the distribution or accumulation of auxin. The polar-localized PIN proteins on the plasma membrane play important roles in plant growth and development as auxin efflux carriers that mediate polar auxin transport (Zegzouti et al., 2006; Zhang et al., 2010). To explore whether RPS6A affects polar auxin transport, by crossing with GFP-fused PIN marker lines, we found that although the fluorescence signals of PIN1-GFP, PIN3-GFP and PIN7-GFP were weakened in *rps6a* roots, there was no change in the polar localization of PIN1-GFP (Figure 4A and B), PIN3-GFP (Figure 4C and D) and PIN7-GFP (Figure 4E and F). While the fluorescence intensity and polar localization of PIN2-Venus did not change (Supplemental Figure 5). To further verify the CSLM results, we performed Western blot assay and found the abundance of PIN1-GFP, PIN3-GFP was decreased in *rps6a* mutant (Figure 4G). Overall, we propose that RPS6A regulates polar auxin transport by modulating the abundance but not the polarity of PIN proteins.

**Figure 4.**
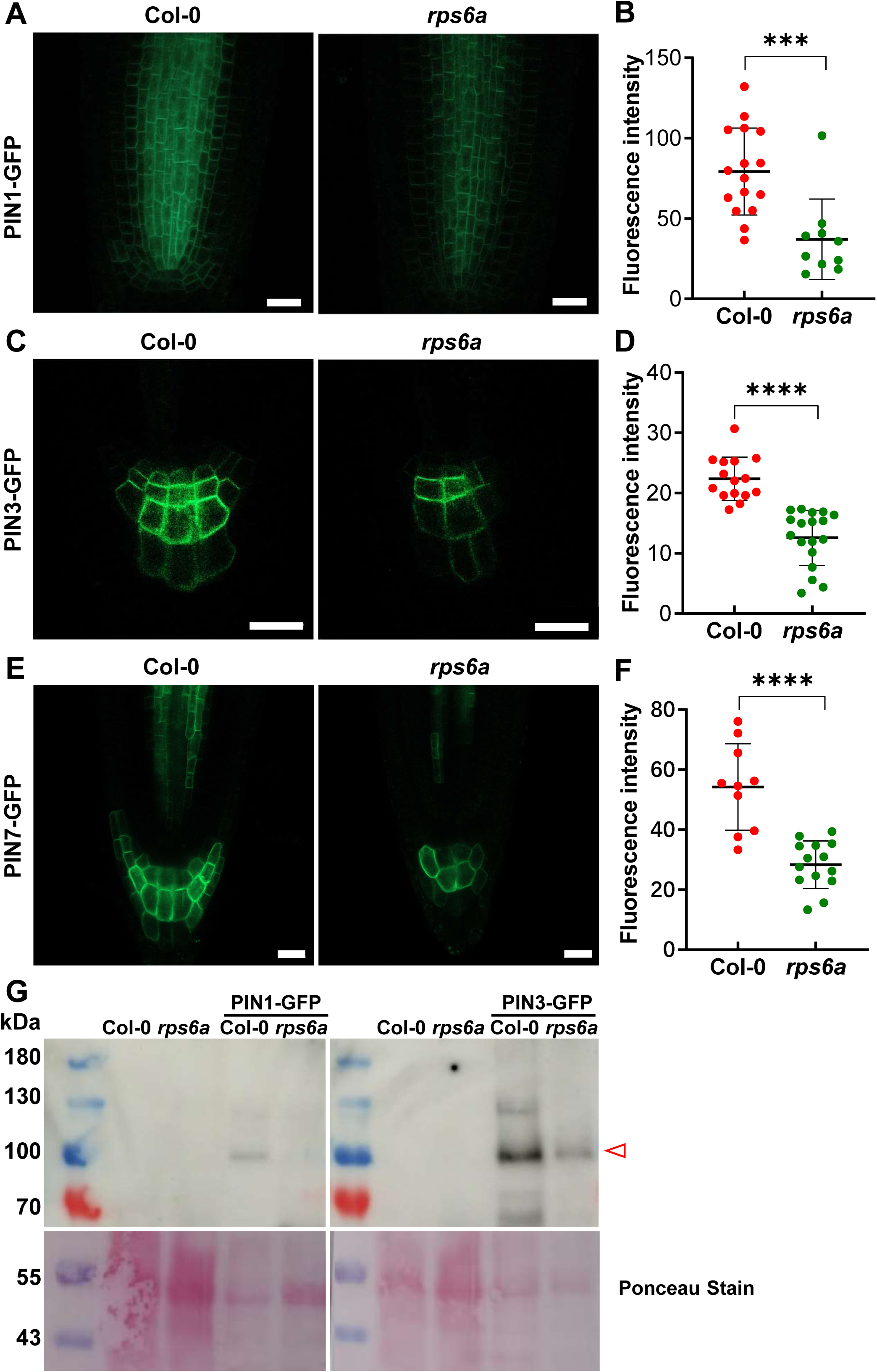
PIN1-GFP, PIN3-GFP and PIN7-GFP fluorescence signals of *rps6a* decreased (**A, C, E**) The *PIN1-GFP*, *PIN3-GFP* and *PIN7-GFP* signals decreased in the *rps6a* mutant. Four-day-old *PIN1-GFP*, *PIN3-GFP* and *PIN7-GFP* Col-0 and *rps6a* seedlings were grown on MS media. The root tips were imaged via confocal laser scanning microscopy (CLSM) in the GFP channel at 40×; scale bar, 20 μm. (**B**, **D, F**) The GFP signal in panels A, C, and F was measured to indicate the fluorescence intensity. n = 16, 10, 15, 18, 10, or 14. The dots represent individual values, and the lines indicate the mean ± SD. \*\*\**p*<0.001, and \*\*\*\**p*<0.0001 according to Welch’s t test. (**G**) Western blot shown the abundance of PIN1-GFP, PIN3-GFP decreased in *rps6a* mutant. The sample was detected by an anti-GFP antibody.

RPS6A is a component of the 40S small ribosome subunit and is involved in protein translation. Therefore, in order to explore whether RPS6A affects the transcription of PIN proteins, we analyzed the expression level of *PIN* genes in *rps6a* and WT by RT-qPCR. The results showed that there was no significant difference in the expression level of *PIN1*, *PIN4*, *PIN6* and *PIN8* in *rps6a* mutant and WT (Supplemental Figure 6). The expression levels of *PIN2* and *PIN5* decreased, and *PIN3* and *PIN7* increased (Supplemental Figure 6). The results of RT-qPCR and RNA-seq are generally consistent. The differences in the transcription and protein levels of *PIN1*, *PIN3* and *PIN7* (Figure 4) in *rps6a* and WT, suggested that RPS6A, as a ribosomal protein, regulates plant growth and development by affecting the polar auxin transport.

### RPS6A regulates root meristem size and cell production

To further investigate the molecular mechanism by which RPS6A is involved in auxin-regulated root growth and development, we performed RNA-seq profiling experiments with WT and *rps6a Arabidopsis* seedlings treated with IAA or the solvent control DMSO. Changes in the expression of auxin-related genes were analysed in WT and *rps6a* plants under IAA treatment compared with those under the control (DMSO) treatment (Supplemental Figure 6). Compared with that in the WT, the number of IAA-induced genes in *rps6a* significantly increased, but the number of overlapping genes upregulated after treatment decreased (Figure 6A and Supplemental Figure 7). We performed GO enrichment analysis with DEGs and found most of the DEGs were inriched in defense-related pathways (Supplemental Figure 8). We draw heatmap with auxin-related genes and found that RPS6A could affect the auxin signalling pathway (Supplemental Figure 9). In addition, we further analyzed RNA-seq data and found that RPS6A can affect the expression of root development related genes in *Arabidopsis* (Supplemental Figure 10). The regulation of RPS6A on the growth and development of roots may be related to the effect of RPS6A on the auxin signaling pathway.

In order to check the effect of RPS6A on auxin signaling, we performed RT-qPCR analysis of some auxin downstream marker genes. In the *rps6a* mutant, the expression levels of *ABP1*, *LBD39*, *IAA31* and *IAA33* decreased and the expression levels of *AFB5*, *GH3.3* and *TIR1* increased. (Figure 6B and C). These results were consistent with our RNA-seq results, indicating that RPS6A affects auxin signal pathway.

Auxin plays a central role in root growth and development by regulating root cell division, growth and differentiation (Saini et al., 2013; Roychoudhry and Kepinski, 2022). In *Arabidopsis*, the stem cell niche and the size of the root meristem are maintained by the quiescent center (QC), stele stem cells, columella stem cells and cortex/endodermis initial cells (Dello Ioio et al., 2007; Yamada et al., 2020). QC cells exhibit low levels of cell division activity, which is essential for root stem cell maintenance. To understand the origin of the root growth defect in *rps6a*, we measured the size of the root meristem of the primary root in WT and *rps6a* seedlings, which was determined by the number of cortex cells between the QC and elongation zones. Compared with those of the WT, the QC zone of *rps6a* was deficient, and the cortex cell number and meristem length were significantly shorter (Figure 5). The *rps6b* mutant presented the same phenotype (Supplemental Figure 11). These observations clearly show that QC function is impaired in the *rps6a* mutant compared with the WT and that the slow root growth of *rps6a* is due to reduced meristem activity.

**Figure 5.**
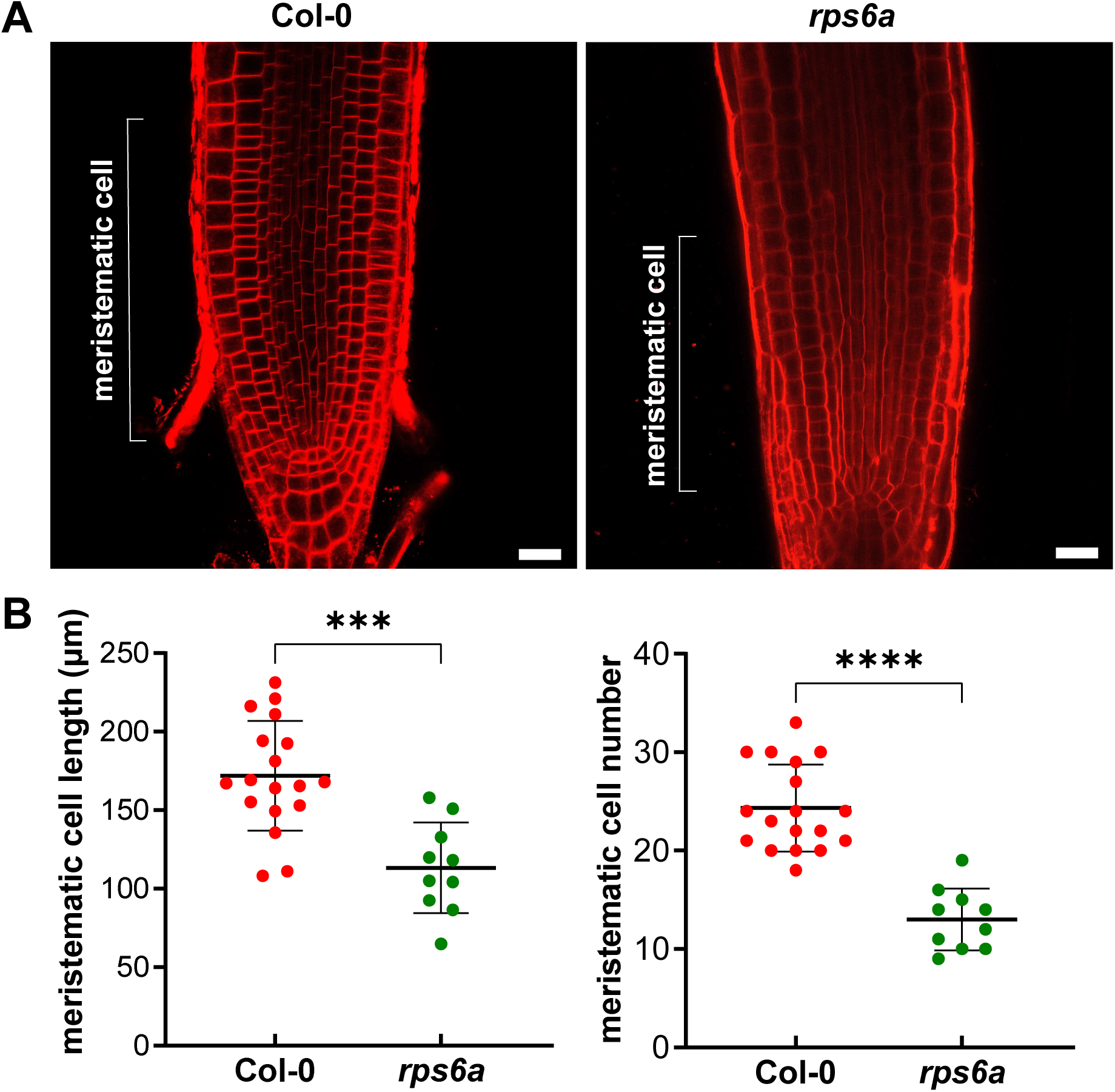
Meristematic cells in the roots of the *rps6a* mutant were reduced. (A) Four-day-old Col-0 and *rps6a* seedlings were stained with FM4-64 for 15 min. Root tips were imaged via confocal laser scanning microscopy (CLSM) for the FM4-64 channel, 40×; scale bar, 20 μm. (B) Quantification of meristematic cell length and cell number in Figure B, n=18, 10. The dots represent individual values, and the lines indicate the mean ± SD. \*\*\*\**p*<0.0001 according to Welch’s t test.

**Figure 6.**
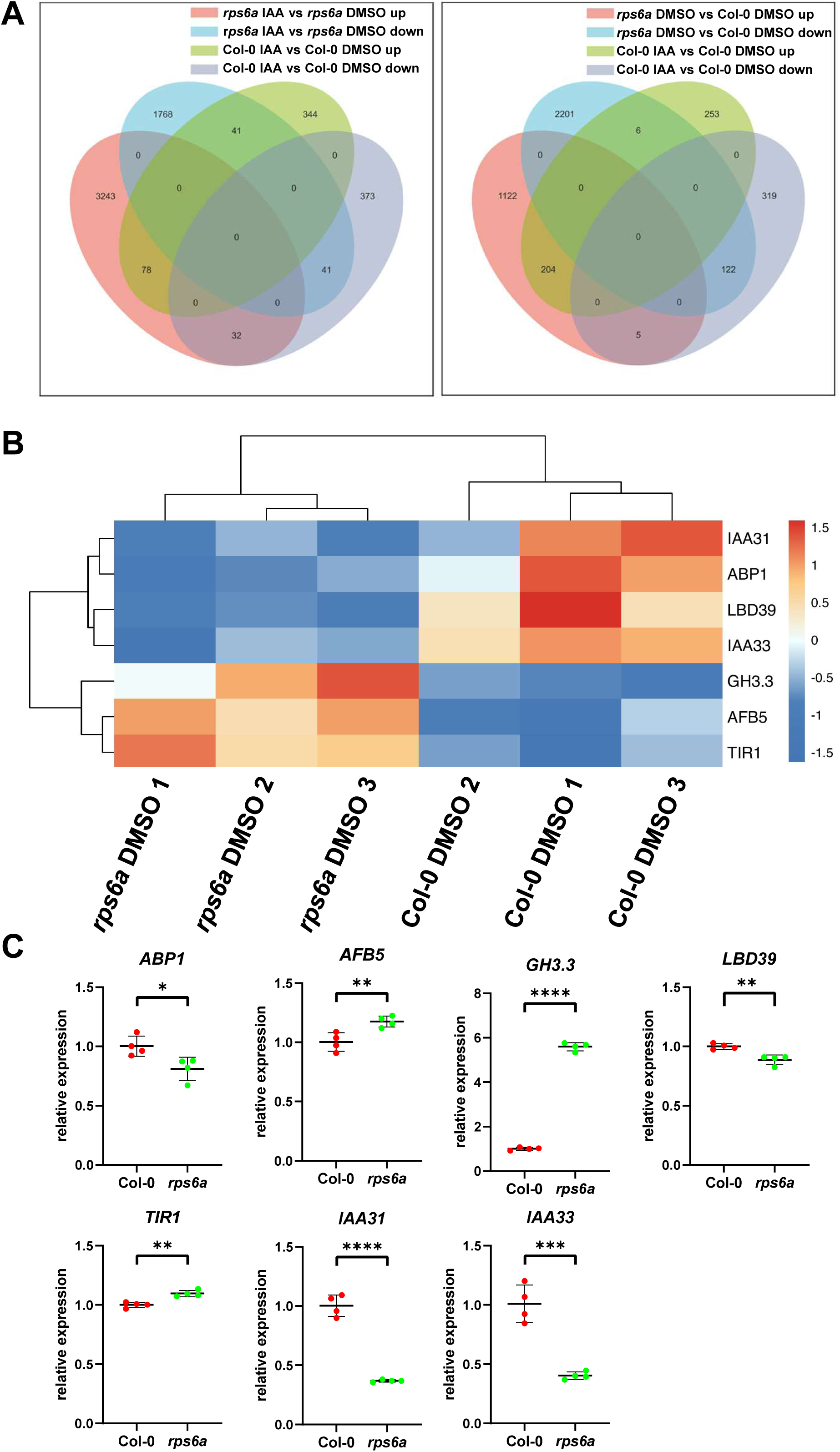
The RNA-seq results revealed that gene expression changed in the mutant after auxin treatment. (A) Venn diagram showing the number of DEGs (FDR< 0.05, versus Col-0) in various pairwise comparisons. (B) Heatmap showing the expression patterns of auxin-related genes *ABP1*, *AFB5*, *GH3.3*, *TIR1*, *LBD39*, *IAA5*, *IAA19*, *IAA31* and *IAA33* in the *rps6a* mutant and Col-0. Seven-day-old Col-0 and *rps6a* seedlings were grown vertically on MS media and treated with 1 µM IAA or DMSO (as the mock control) for 3 h. Hierarchical clustering of log_2_FC values (Col-0 vs *rps6a*) was calculated via DESeq2 analysis. (C) RT-qPCR analysis show relative expression levels of auxin-related genes *ABP1*, *AFB5*, *GH3.3*, *TIR1*, *LBD39*, *IAA5*, *IAA19*, *IAA31* and *IAA33*. Dots represent individual values (Col-0 is red dots and *rps6a* is green dots), and lines indicate mean ± SD. \**p*<0.05, \*\**p*< 0.01, \*\*\**p*<0.001 and \*\*\*\**p*<0.0001 according to Welch’s t test.

## Discussion

The ribosome plays a central role in protein translation. In this study, we performed phenotypic analysis, cell biological experiments and transcriptome profiling to investigate the role of *RPS6* in plant growth and development. The *rps6a* mutant presented shortened primary roots, fewer lateral roots, defective cotyledon veins and fewer root meristem cells, which are all auxin-deficient phenotypes. We did not do much analysis of the *rps6b* CRISPR mutant due to its similar phenotypes as *rps6a.* Notably, *rps6b* and *rps6a* have similar phenotypes in primary root length, lateral root number, and root meristem cell number. Thus, according to the above results and previous studies, we believe that *RPS6A* and *RPS6B* are functionally redundant and interchangeable (Creff et al., 2010).

The DR5 signal revealed that the auxin concentration was reduced in the tip of the *rps6a* mutant. Notably, the *rps6a* mutant was relatively insensitive to auxin treatment, suggesting a defect in auxin signalling. Further subcellular localization observations revealed that the abundance of the PIN1, PIN3 and PIN7 proteins was lower than that of Col-0. When we observed the DR5rev::GFP signal, the morphology of the QC region was abnormal in the *rps6a* mutant, and a double layer of QC cells appeared. This may be due to the lower auxin concentration in the root tip.

The process of protein expression involves many regulatory processes, such as transcriptional regulation, posttranscriptional regulation, and translational regulation. In most cases, scientists tend to focus on RNA-seq results to infer gene expression levels. However, only approximately 40% of mRNA abundances are clearly correlated with their translated protein abundances (Vogel and Marcotte, 2012). Different protein or rRNA compositions of the ribosome can affect the translation of specific mRNAs (Dalla Venezia et al., 2019). Here, we found that the loss of *rps6a* resulted in a decrease in the abundance of the auxin transporters PIN1, PIN3, and PIN7, which led to a decreased auxin concentration and an auxin-deficient phenotype. In the RNA-seq results, the transcript levels of *PIN3* and *PIN7* were changed compared with wild type. These results suggested that RPS6A might affect plant growth and development by affecting protein translation and transcription, which is in line with its biochemical functions.

## Materials and Methods

### Plant Materials and Growth Conditions

The *Arabidopsis thaliana* (L.) mutants and transgenic lines were all of the Columbia-0 ecotype (Col-0) background. The marker lines *DR5rev::GFP* (Friml et al., 2003), *pPIN1::PIN1-GFP* (Xu et al., 2006), *pPIN3::PIN3-GFP* (Zádníková et al., 2010) and *pPIN7::PIN7-GFP* (Kleine-Vehn et al., 2010) were published previously. The mutant *rps6a* (SALK_048825C) in the Columbia-0 (Col-0) background was also reported previously, and Col-0 was used as a control.

For phenotyping of seedlings or pharmacological experiments, surface-sterilized seeds were sown on solid Murashige and Skoog (MS) media [0.5× MS media supplemented with 1% (w/v) sucrose, 0.8% (w/v) phytoagar, MES buffer, pH 5.9], stratified at 4°C for 2 d, and then grown vertically in a growth chamber at 21°C with a 16-h light/8-h dark photoperiod.

### Pharmacological treatments

For auxin and its analogue treatment assays, *Arabidopsis* seeds were sown on vertical plates with MS media. After 2 d of stratification, the plants were transferred to a growth chamber as described in the “Plant material and growth conditions” session for 4 or 7 d. Then, the seedlings were placed on MS media supplemented with the indicated chemicals, including indole 3-acetic acid (Sigma, I2886), 2,4-dichlorophenoxyacetic acid (Solarbio, D8100), α-naphthaleneacetic acid (Solarbio, N8010) and dimethyl sulfoxide (DMSO, Sigma, D4540), which served as mock controls. The root length and lateral root number were measured after an additional 3 d of growth.

For long-term hygromycin B (Hyg) treatment assays, *Arabidopsis* seeds were sown on vertical plates with MS media supplemented with the indicated chemicals, including Hyg (BBI, A600230-0001) and DMSO as the mock control. After 2 d of stratification, the plates were transferred to a growth chamber as described in the “Plant material and growth conditions” section. Seedlings were observed via confocal laser scanning microscopy (CLSM) after they had grown for 4 d, or primary root length was measured after they had grown for 7 d.

### Imaging by confocal laser scanning microscopy (CLSM)

Fluorescence imaging was performed via a Zeiss LSM980 confocal laser scanning microscope with a GaAsP detector (Zeiss, Germany). The manufacturer’s default settings (smart mode) were used for imaging proteins tagged with GFP (excitation, 488 nm; emission, 495–545 nm) or FM4-64 (excitation, 543 nm; emission, 600–700 nm). All of the images were obtained at 8 bit depth with 2× line averaging. The images were analysed and visualized with the Fiji program.

For the IAA treatment assays used to observe the DR5rev::GFP signal, 3-day-old seedlings were transferred to MS media supplemented with 3-acetic acid or DMSO as the mock control and observed after 24 hours.

To image FM4-64-stained cells, 4-day-old seedlings were incubated in FM4-64 [N-(3-triethylammoniumpropyl)-4-(6-(4-(diethylamino) phenyl)hexatrienyl)pyridinium dibromide, Invitrogen, T13320] (1 μg/ml in liquid MS medium) for 15 minutes and rinsed three times in liquid MS medium before observation.

### Root gravitropism assay

Four-day-old seedlings were arranged on MS media and rotated 90°. The plants were grown as described in the “Plant material and growth conditions” session and were observed every two hours.

### Image analysis and morphological analysis

For observation of the seedling root phenotype, photographs were taken via a camera (Sony A6000 with a macrolens), and then, the primary root length or root tip angles were analysed with ImageJ (Schindelin, et al., 2012). Lateral root numbers were counted directly. The veins of the cotyledons were observed via a Nikon SMZ18. To observe the fluorescence in the roots, all the images were obtained via CLSM, and the fluorescence intensity was measured via ImageJ.

### RNA-seq analysis

For RNA-seq analysis, 7-day-old Col-0 and *rps6a* seedlings were treated with 1 µM IAA or DMSO for 3 h. Every group had three independent 0.1 g samples. The samples were frozen in liquid nitrogen and sent to the Beijing Genomics institution for RNA-seq. Lists of DEGs are shown in Supplemental Tables 1 to 4. List of auxin and root-related genes are shown in Supplemental Tables 5 and 6.

### RT-qPCR

Quantitative PCR with reverse transcription (RT-qPCR) was used to examine the transcripts of *rps6a* mutants and Col-0, *ACTIN7* (*AT5G09810*) was used as an internal reference. Total RNA was extracted by FastPure Universal Plant Total RNA Isolation Kit (Vazyme). Then 1 ug of the RNA sample was used for reverse transcription by 5×PeimeScript RT Master Mix. The cDNA of corresponding genes and *ACTIN7* was analysed using the SYBR Premix Ex Taq (Takara) with a Bio-Rad CFX Connect Real-Time System. Relative transcript level of the examined genes was normalized to the expression of *ACTIN7*, and was calculated by setting WT or a certain tissue as 1, and the data are presented as mean ±SD from three biological replicates. The primers used are shown in supplemental tables.

### Protein extraction and Western blot

Col-0, *rps6a*, *PIN1::PIN1-GFP*, *rps6a PIN1::PIN1-GFP*, *PIN3::PIN3-GFP*, *rps6a PIN3::PIN3-GFP* seedlings (7d) were used for total protein extraction. After harvesting, the samples were ground in liquid nitrogen, resuspended in 10% TCA acetone solution and stored in −20L for 3h. Then centrifuge and resuspend the precipitate with acetone, repeat five times to completely clean precipitate. Finally resuspend the precipitate with protein extraction buffer (1×PBS, 9M urea, 10mM DTT) and spun down to discard the cell debris.(Koontz, 2014) The protein samples were then analyzed by western blot. PIN1-GFP, PIN3-GFP was detected by an anti-GFP antibody (JL8, Clontech, 1:2000). HRP activity was detected by the HRP-labeled Goat Anti-Mouse IgG(H+L) (Beyotime) and imaged with AMERSHAM ImageQuant 800.

### Dark treatment

*Arabidopsis* seeds were sown on vertical plates with MS media. After 2 d of stratification, the plates were transferred to a growth chamber as described in the “Plant material and growth conditions” session for 1 d and then wrapped in tin foil for 3 d. The hypocotyl length was measured after 4 d.

## Quantification and Statistical Analysis

Most experiments were repeated at least three times independently, with similar results obtained. For measurements of primary root length and root tip angles, photographs or scans were analysed with the ImageJ program (https://imagej.nih.gov/ij/download.html). The fluorescence intensity of the CLSM images was quantified with Fiji (https://fiji.sc/)(Schindelin)(Schindelin et al., 2012). Data visualization and statistical analysis were mostly performed with GraphPad Prism 8. For the bending curves of the root tips, polar graphs were generated via Origin 2023, n and p values are indicated in the figures or legends.

## SUPPLEMENTAL INFORMATION

Supplemental information is available at xx Online.

## Supplemental Figure legends

**Supplemental Figure 1.**
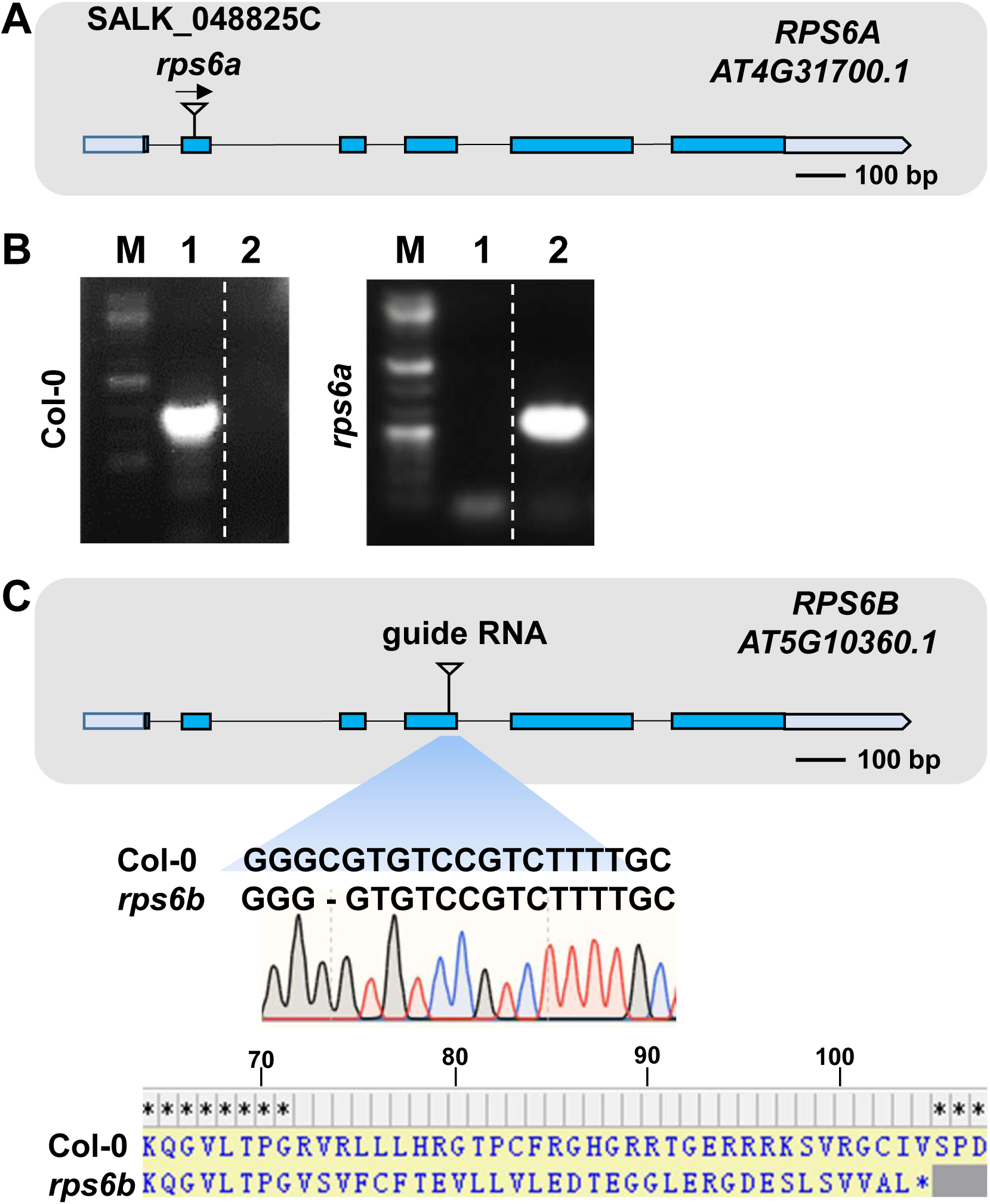
Genotyping of the *rps6a* and *rps6b* mutants. (A) T-DNA insertion information for the *rps6a* mutant. The blue boxes indicate exons, the black lines represent introns, the gray boxes on both sides indicate noncoding (UTR) regions, and the black arrows indicate the direction of T-DNA insertion. Scale bar, 200 bp. (B) PCR results of *rps6a* mutation identification. 1, LP+RP; 2, LBb1.3+RP; Col-0 was used as a control. (C) CRISPR information for the *rps6b* mutant. The guide RNA is located on the third exon. The absence of the single base C leads to differences in the amino acid chain and premature termination.

**Supplemental Figure 2.**
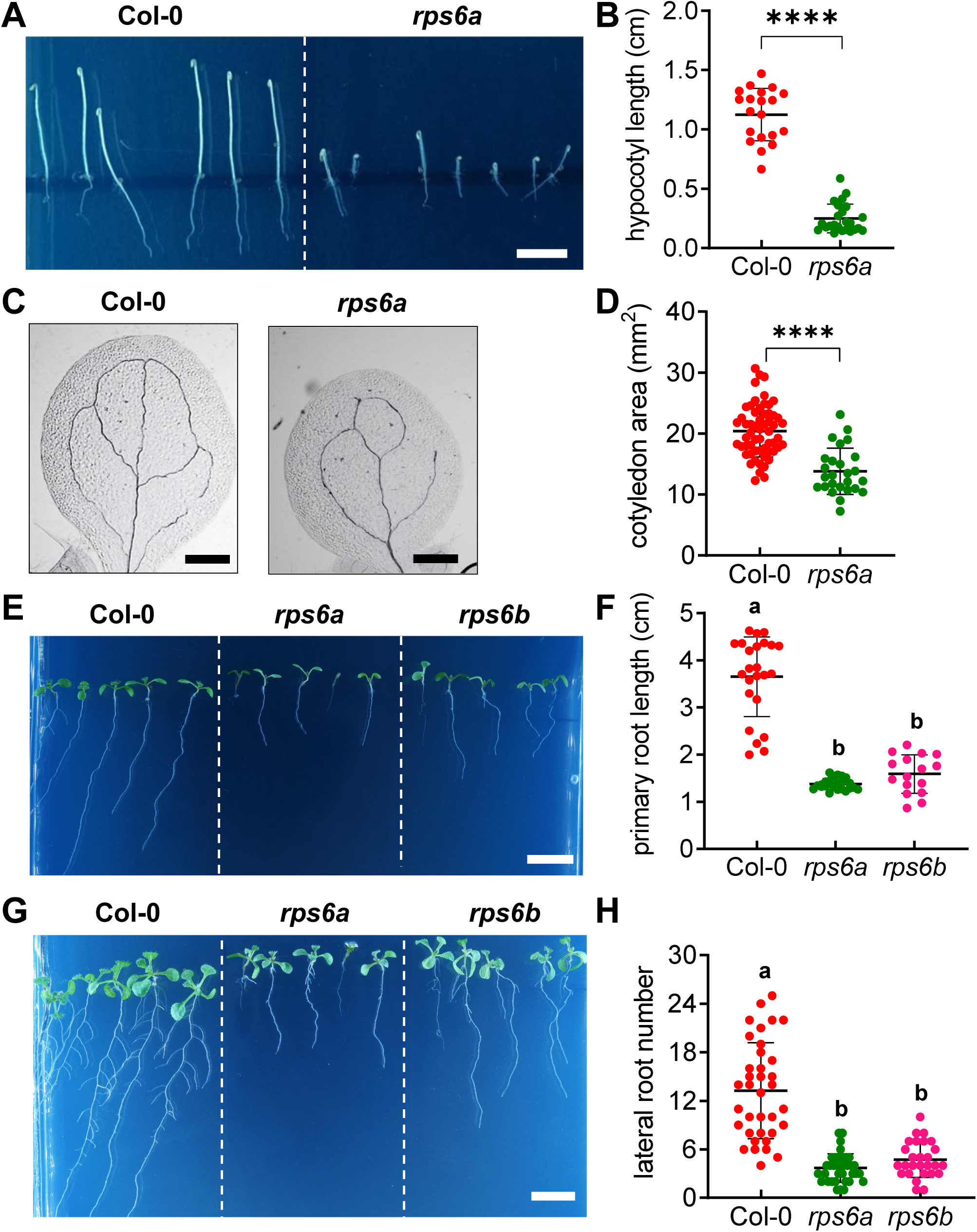
The hypocotyls and cotyledons of *rps6a* are defective. (A) The *rps6a* mutant has a shorter hypocotyl. Four-day-old seedlings were grown on MS media in darkness. Scale bar, 1 cm. (B) Quantification of hypocotyl lengths in Figure A, n=20,24. The dots represent individual values, and the lines indicate the mean± SD. \*\*\*\**p*<0.0001 according to Welch’s t test. (C) The *rps6a* mutant has obvious defects in cotyledon development. Compared with those of Col-0 cotyledons, the cotyledons of *rps6a* cotyledons are smaller and have at most two intact rings and broken veins. Scale bar, 1 cm. (D) Quantitation of the cotyledon area shown in Figure C, n=58,26. The dots represent individual values, and the lines indicate the mean± SD. \*\*\*\**p*<0.0001 according to Welch’s t test. (**E, G**) Both the *rps6a* and *rps6b* mutants presented shorter primary roots and fewer lateral roots. Seven-day-old seedlings in Figure E and 10-day-old seedlings in Figure G were grown on MS media. Scale bar, 1 cm. (**F, H**) Quantification of the primary root lengths in Figure E and the number of lateral roots in Figure G, n=24, 24, 16. The dots represent individual values, and the lines indicate the mean± SD. \*\*\*\**p*<0.0001 according to Welch’s t test.

**Supplemental Figure 3.**
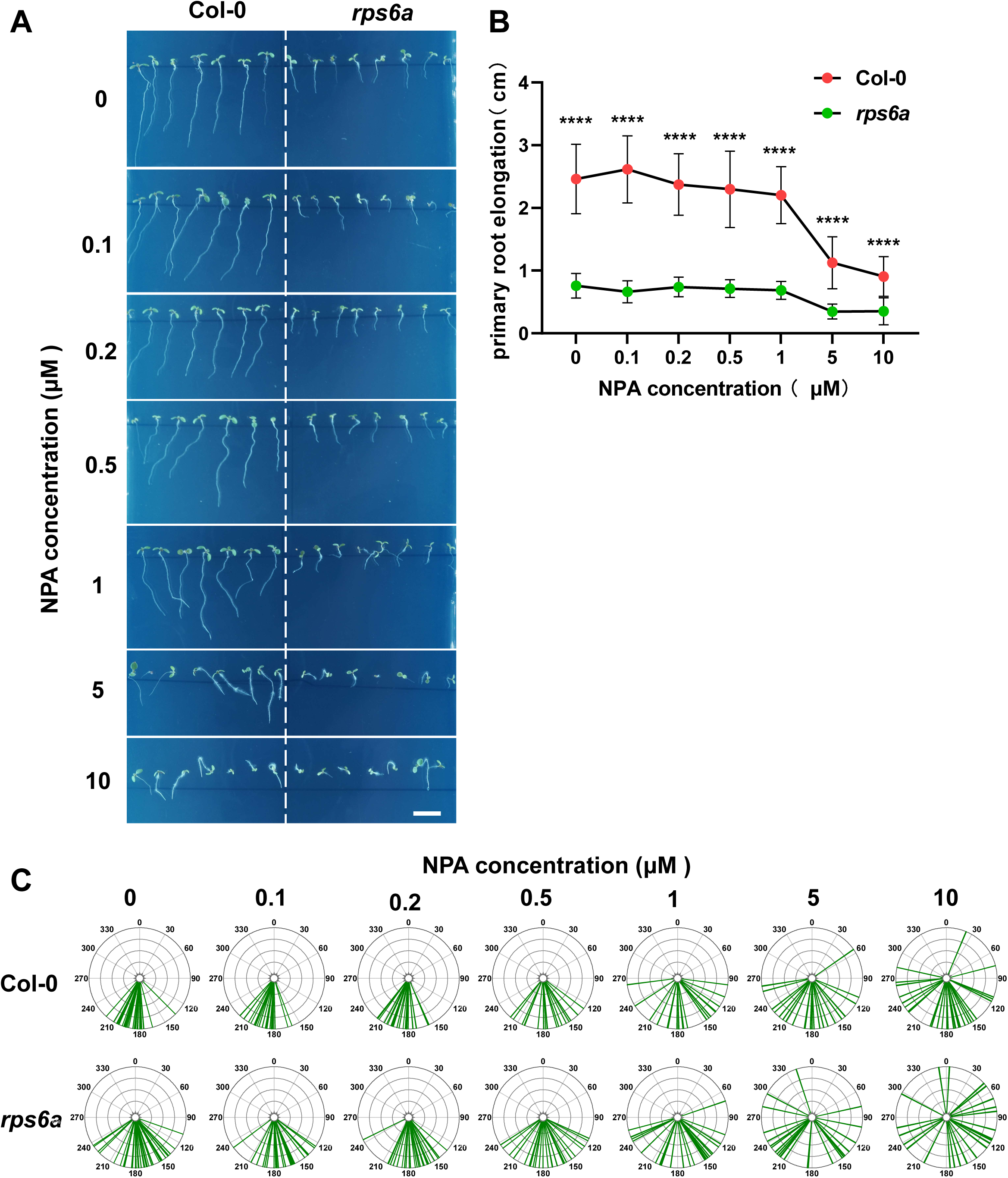
NPA treatment resulted in a more severe gravitrotropic defect in *rps6a*. (A) Col-0 and *rps6a* were treated with 0, 0.1, 0.2, 0.5, 1, 5, 10 μM NPA. Seven-day seedlings, Scale bar, 1 cm. (B) Quantitation of primary root lengths in figure A. Col-0, n =32, 20, 26, 20, 23, 22, 30. *rps6a*, n = 41, 25, 28, 25, 27, 25, 21. The dot represent average value, and the lines indicate the ± SD. \*\*\*\**p*<0.0001 according to Welch’s t test. (C) Root tip angle of Figure A. The root tip angle of *rps6a* was more dispersed and the phenotype is more pronounced after NPA treatment.

**Supplemental Figure 4.**
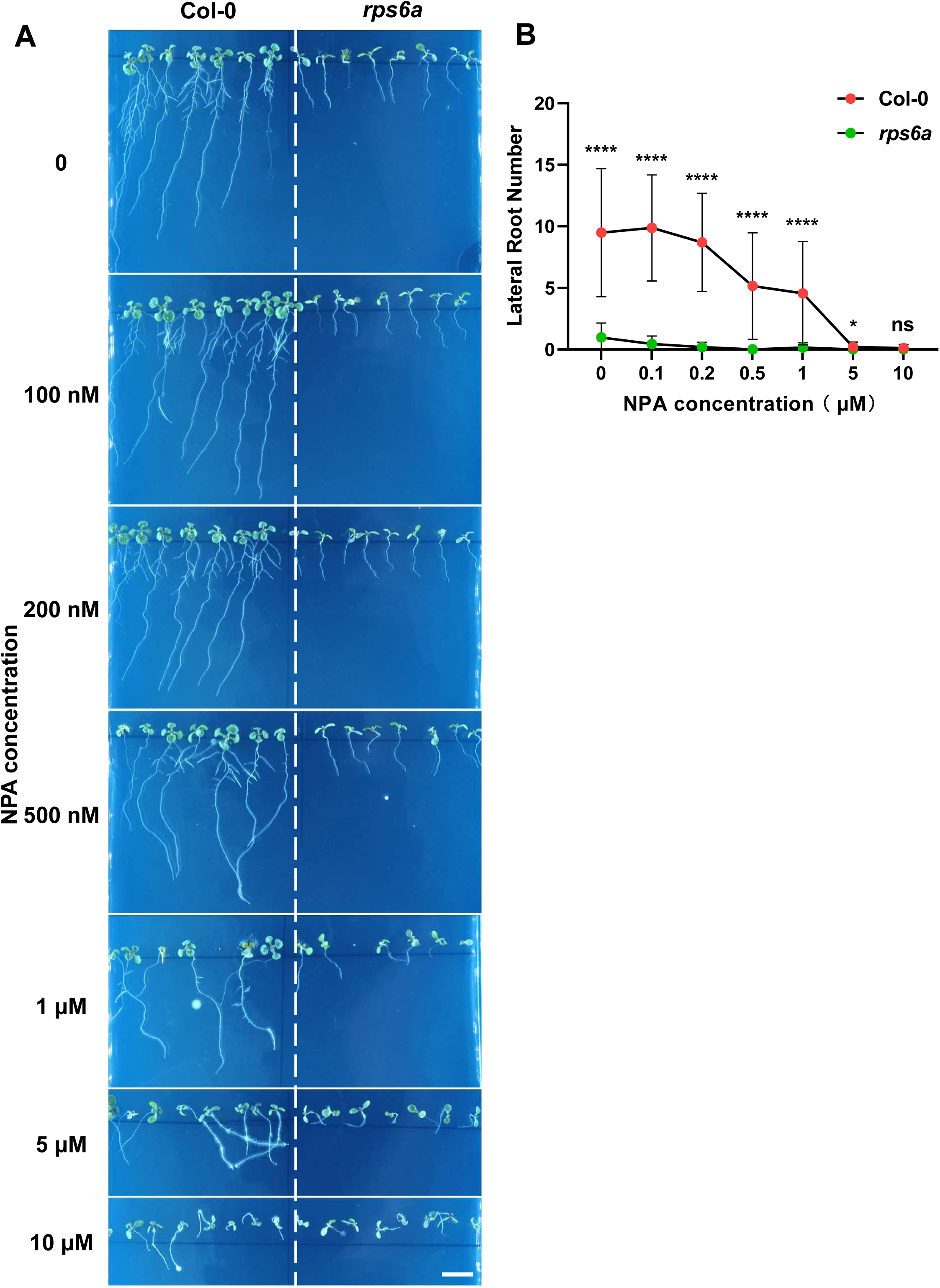
Lateral root number of *rps6a* was sensitive to NPA treatment. (A) Treat Col-0 and *rps6a* with 0, 0.1, 0.2, 0.5, 1, 5, 10 μM NPA. Ten-day-old seedlings, Scale bar, 1 cm. (B) Quantitation of primary root lengths in figure A. Col-0, n =39, 23, 26, 20, 23, 25, 30. *rps6a*, n = 41, 25, 27, 26, 25, 25, 25.The dot represent average value, and the lines indicate the ± SD. \**p*<0.05, \*\*\*\**p*<0.0001 according to Welch’s t test.

**Supplemental Figure 5.**
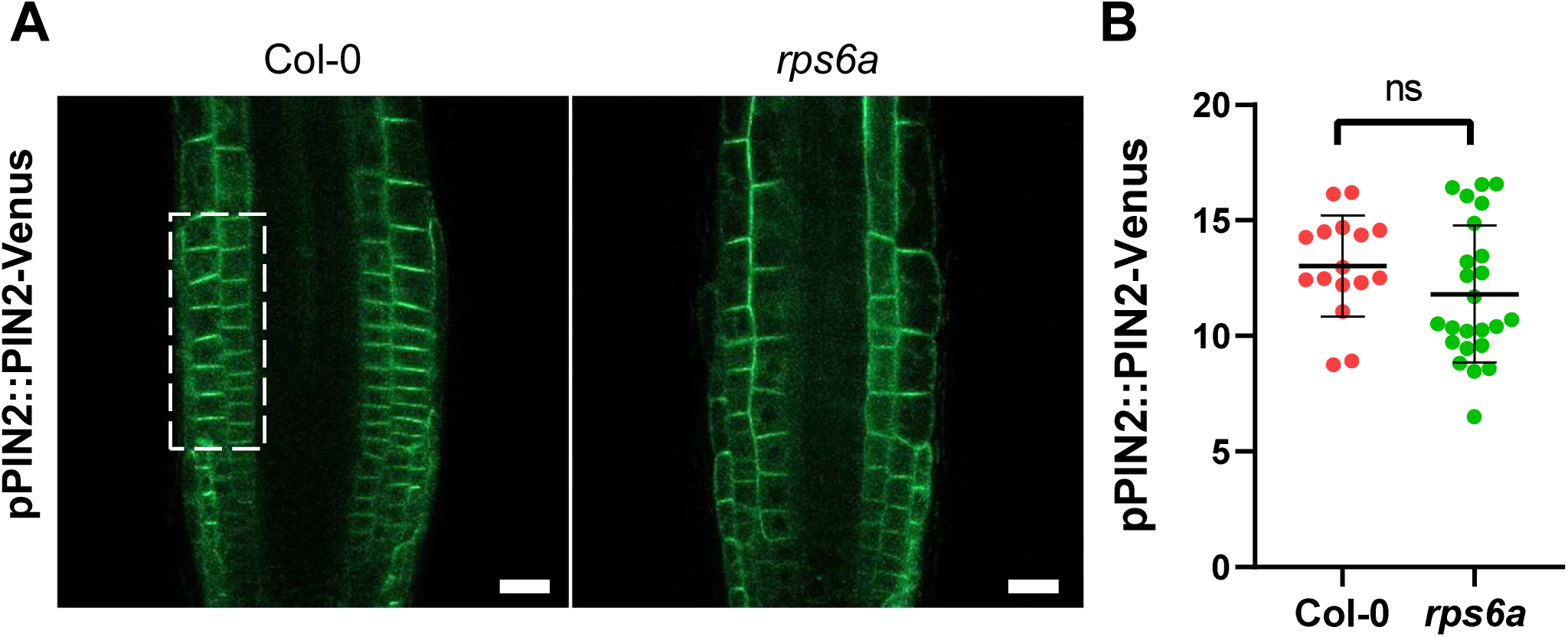
PIN2-Venus fluorescence signals of *rps6a* did not change (A) The *PIN2-Venus* signals did not change in the *rps6a* mutant. Four-day-old *PIN2-Venus* Col-0 and *rps6a* seedlings were grown on MS media. The root tips were imaged via confocal laser scanning microscopy (CLSM) in the GFP channel at 40×; scale bar, 20 μm. (B) The GFP signal in panels A was measured to indicate the fluorescence intensity. n = 16 and 24. The dots represent individual values, and the lines indicate the mean ± SD according to Welch’s t test.

**Supplemental Figure 6.**
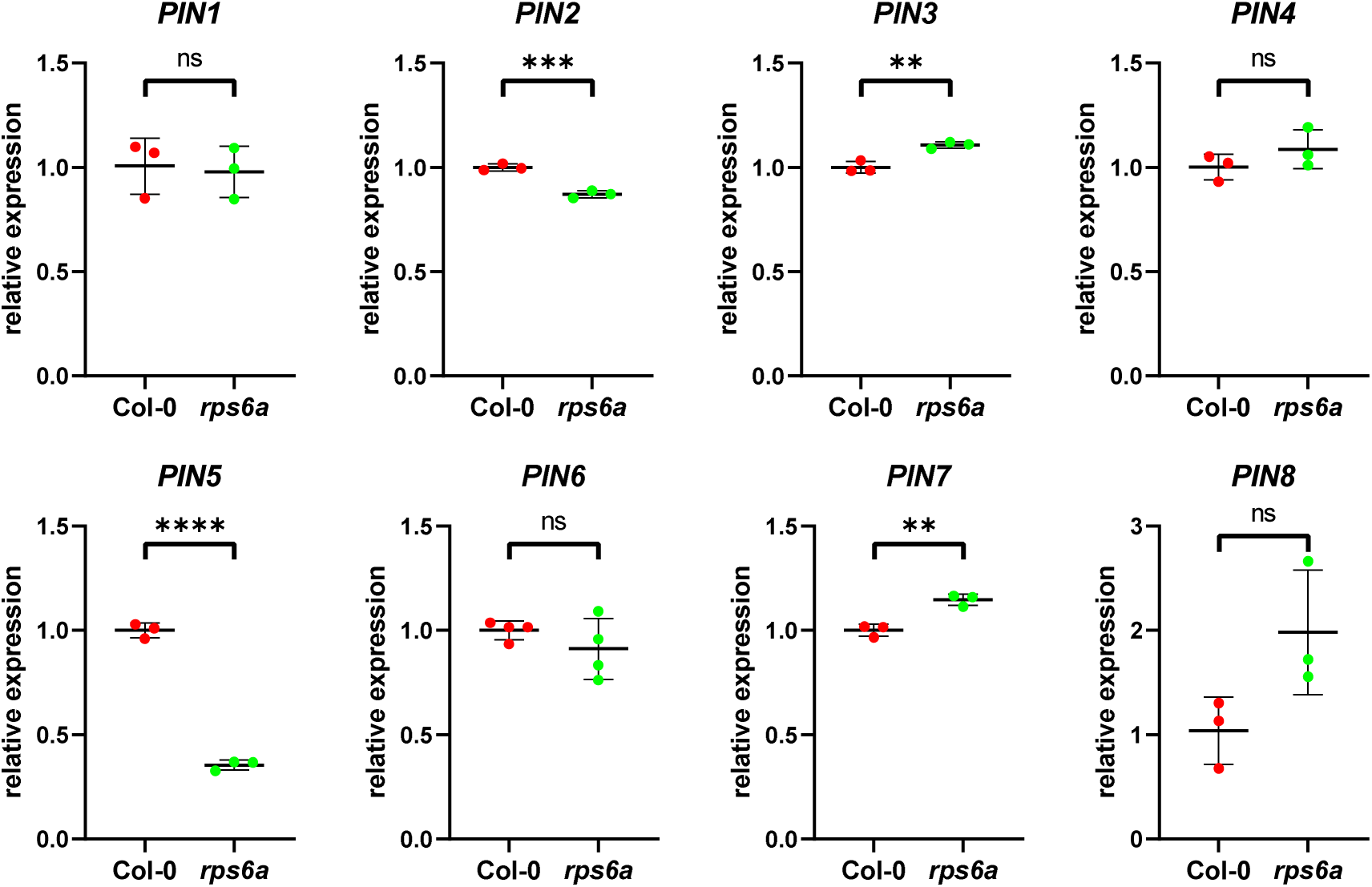
RPS6A affects the expression of auxin related genes RT-qPCR analysis show relative expression levels of *PIN1-PIN8*. Dots represent individual values (Col-0 is red dots and *rps6a* is green dots), and lines indicate mean ± SD. \*\**p* < 0.01, \*\*\**p*<0.001 and \*\*\*\**p*<0.0001 according to Welch’s t test

**Supplemental Figure 7.**
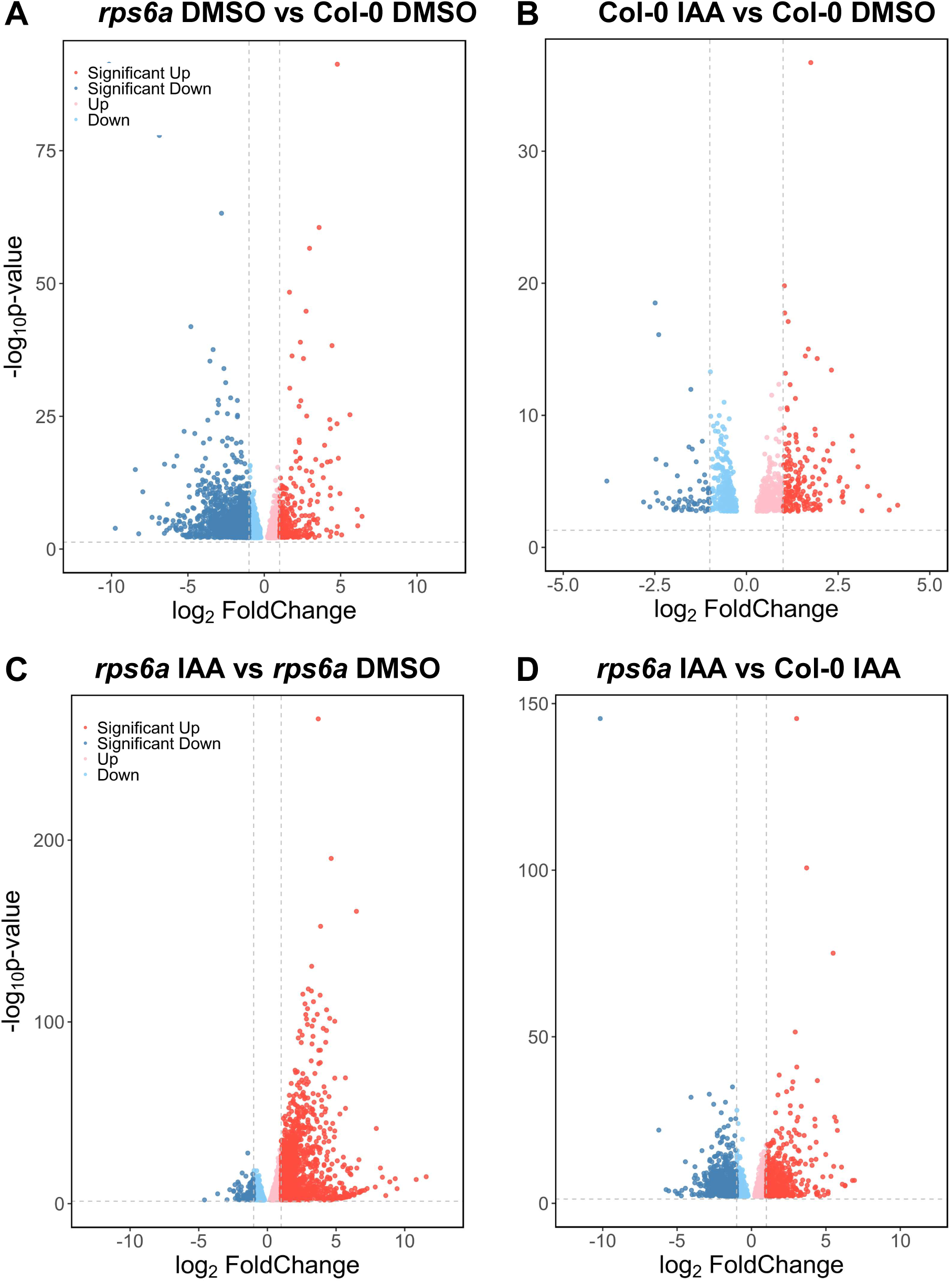
Volcano map of the *rps6a* mutant and Col-0 following IAA treatment. (**A-D**) Volcano map of a schematic representation of the RNA-seq data of 7-day-old Col-0 and *rps6a* seedlings grown vertically on MS media and treated with 1 µM IAA or DMSO (as the mock control) for 3 h. The abscissa is log_2_FC, and the ordinate is -log10 (*p* value). Each point represents a gene (t test: *p*<0.05; log_2_FC>0 or <0 are highlighted with red or blue points). (A) Volcano map of *rps6a* and Col-0 seedlings treated with DMSO. (B) Volcano map of Col-0 seedlings treated with IAA and DMSO. (C) Volcano map of *rps6a* seedlings treated with IAA or DMSO. (D) Volcano map of *rps6a* and Col-0 seedlings treated with IAA.

**Supplemental Figure 8.**
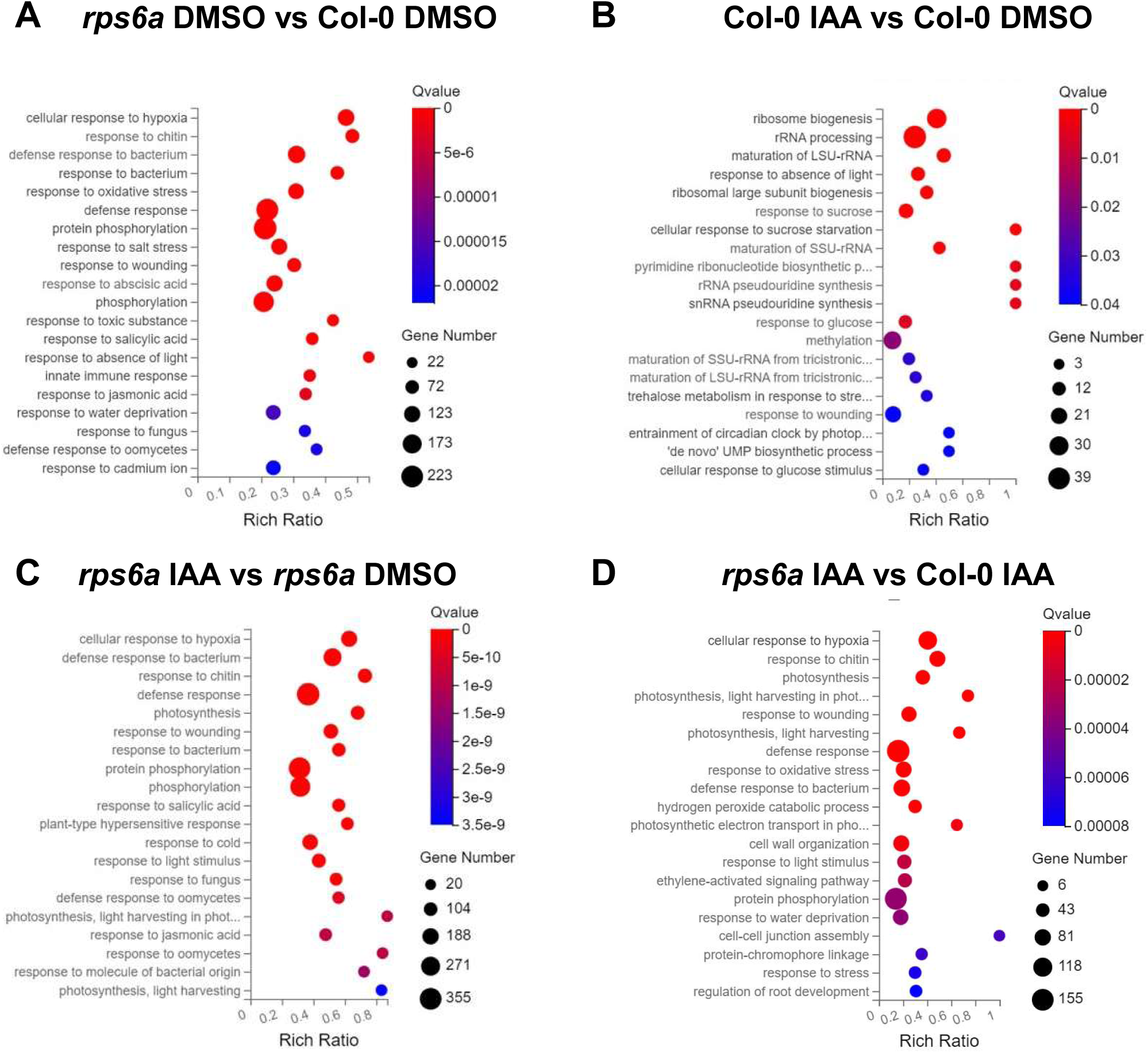
GO inrichment analysis of the *rps6a* mutant and Col-0 following IAA treatment. (**A-D**) GO inrichment analysis of DEGs in Figure 7 (**A**) GO inrichment analysis of *rps6a* and Col-0 seedlings treated with DMSO. (**B**) GO inrichment analysis of Col-0 seedlings treated with IAA and DMSO. (**C**) GO inrichment analysis of *rps6a* seedlings treated with IAA or DMSO. (**D**) GO inrichment analysis of *rps6a* and Col-0 seedlings treated with IAA.

**Supplemental Figure 9.**
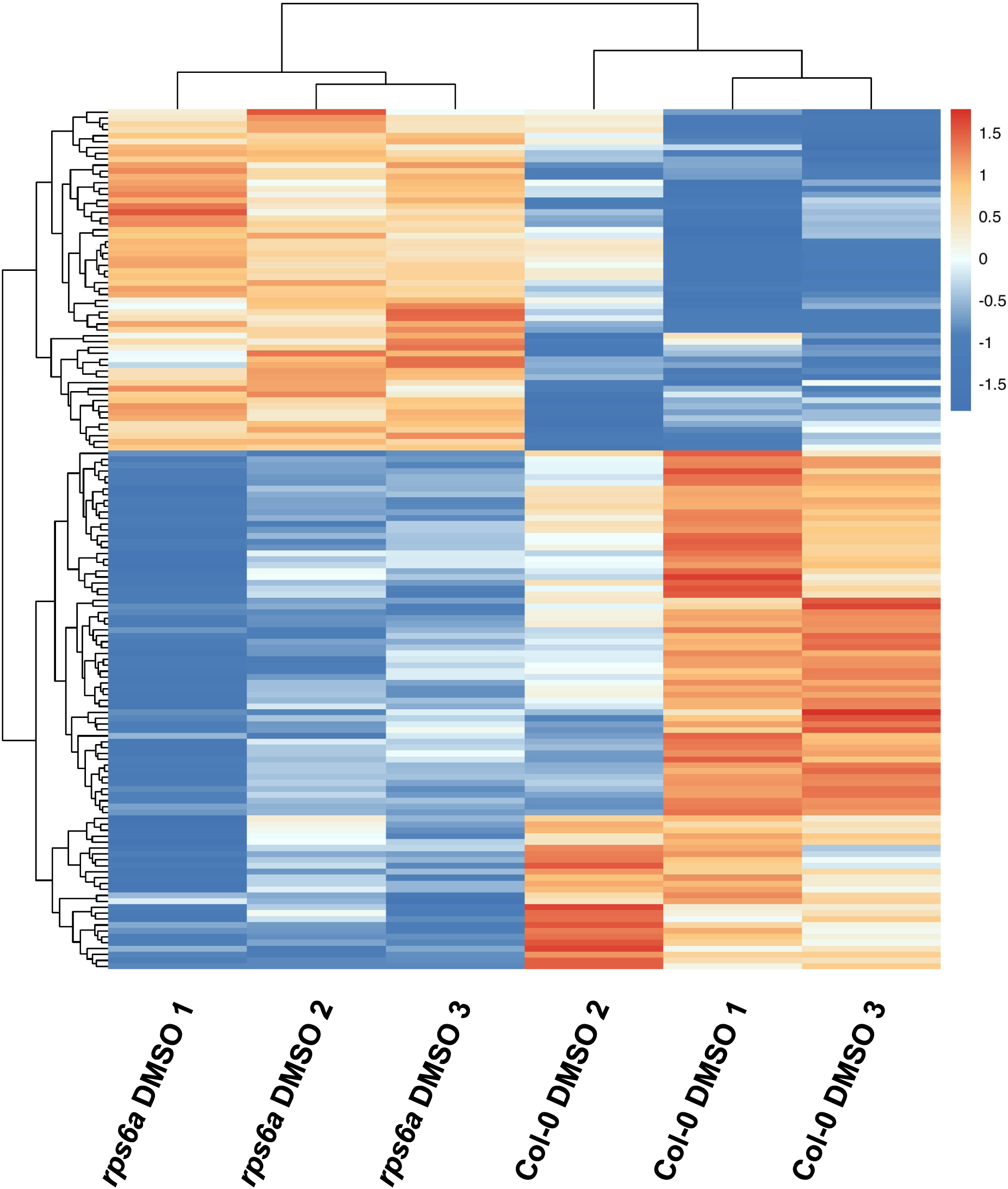
Heatmap of auxin-related genes in the *rps6a* mutant and Col-0 following IAA treatment. Heatmap showing representative auxin-related genes (Supplemental Table 5) expression patterns in the *rps6a* mutant and Col-0 in response to IAA treatment. Seven-day-old Col-0 and *rps6a* seedlings were grown vertically on MS media and treated with 1 µM IAA or DMSO (as the mock control) for 3 h. Hierarchical clustering of log_2_FC values (Col-0 vs *rps6a*) was calculated via DESeq2 analysis.

**Supplemental Figure 10.**
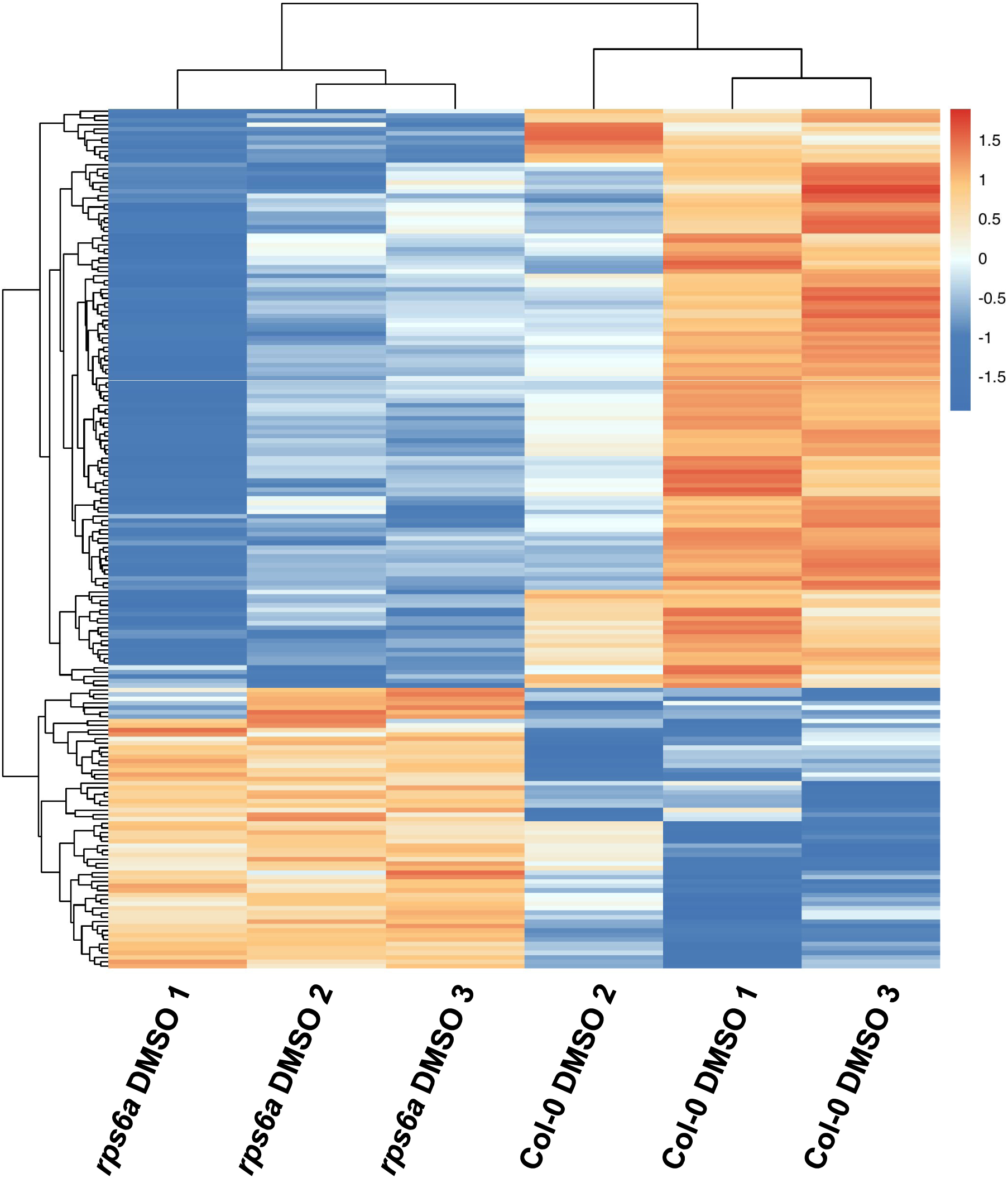
Heatmap of root-related genes in the *rps6a* mutant and Col-0 Heatmap showing all root-related genes (Supplemental Table 6) expression patterns in the RNA-seq result of Col-0 and *rps6a*, revealed that the *rps6a* mutant affected root development. Red indicates upregulated gene expression, while blue indicates downregulated gene expression.

**Supplemental Figure 11.**
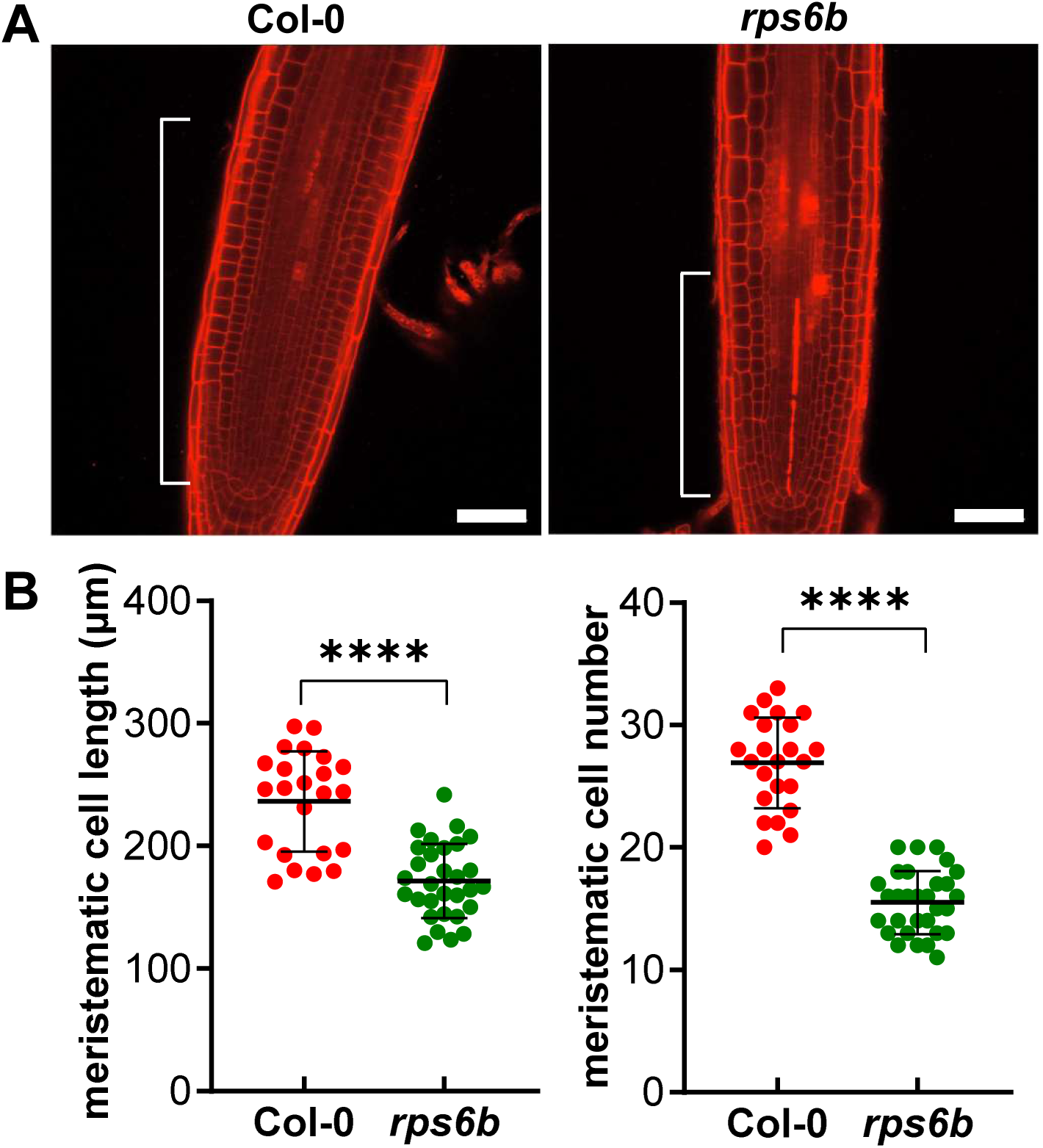
Meristematic cells in the roots of the *rps6b* mutant were reduced. (A) Four-day-old Col-0 and *rps6b* seedlings were stained with FM4-64 for 15 min. Root tips were imaged via confocal laser scanning microscopy (CLSM) for the FM4-64 channel at 40×; scale bar, 20 μm. (B) Quantification of meristematic cell length and cell number in Figure A, n=23, 30. The dots represent individual values, and the lines indicate the mean ± SD. \*\*\*\**p*<0.0001 according to Welch’s t test.

